# A globally influential area-condition metric is a poor proxy for invertebrate biodiversity

**DOI:** 10.1101/2024.10.02.616290

**Authors:** Natalie E. Duffus, Thomas B. Atkins, Sophus O.S.E. zu Ermgassen, Richard Grenyer, Joseph W. Bull, Dan A. Castell, Ben Stone, Niamh Tooher, E.J. Milner-Gulland, Owen T. Lewis

## Abstract

There is increasing demand for standardised, easy-to-use metrics to assess progress towards achieving biodiversity targets and the effectiveness of ecological compensation schemes. Biodiversity metrics based on combining habitat area and habitat condition scores are proliferating rapidly, but there is limited evidence on how they relate to ecological outcomes. Here, we test the relationship between the statutory biodiversity metric used for Biodiversity Net Gain (BNG) in England — and as the basis for new biodiversity credit systems around the world — and invertebrate richness, abundance, and community composition. We find that the combined area-condition BNG metric does not capture the value of arable farmland and grassland sites for invertebrate biodiversity: invertebrate communities were highly variable across sites that had the same type and condition under the BNG metric. We found no reliable relationship between scores under the metric and either invertebrate abundance or species richness, with the risk of the metric undervaluing sites of high invertebrate value. Our results highlight the need to incorporate factors beyond habitat type and condition into site evaluations, and to complement metric use with species-based surveys.

## Introduction

Increased conservation efforts are required to prevent further declines in biodiversity (Díaz et al., 2019; Leclère et al., 2020). Alongside efforts to promote nature recovery, there has been a proliferation of ecological compensation initiatives seeking to enable infrastructure development while compensating for ecological harms (zu Ermgassen et al., 2019). These range from regulations mandating a No Net Loss (NNL) or Net Gain in biodiversity, to voluntary commitments to improve biodiversity by purchasing biodiversity credits (Wunder et al., 2024; zu Ermgassen et al., 2019). These regulations and commitments require robust biodiversity metrics to measure baseline biodiversity value, determine the level of compensation required, and evidence that positive outcomes for biodiversity have been achieved.

Biodiversity metrics based on combining the area, conservation value, and relative condition of habitats (area-condition metrics) are proliferating and have the advantage of being relatively simple to calculate (E. Marshall et al., 2020; Wunder et al., 2024). Such metrics are therefore becoming fundamental to how nature markets measure biodiversity. However, unless these metrics are reliable proxies of wider biodiversity, policy and nature markets using them will fail to improve, or may even be detrimental to, the nature they aim to benefit. Despite widespread concern that area-condition metrics may not provide good proxies for biodiversity, risking sub-optimal ecological outcomes (Duffus et al., 2024; Hanford et al., 2017; Hawkins et al., 2022; C. A. M. Marshall et al., 2024; Rampling et al., 2023; Simpson et al., 2022; Thorn et al., 2018; zu Ermgassen et al., 2023), studies investigating the relationship between area-condition metric scores and community assemblages assessed using primary data and across multiple taxa are lacking.

In England, an area-condition metric – the Statutory Biodiversity Metric – has been adopted to support a requirement for new housing and infrastructure projects to deliver a 10% Biodiversity Net Gain (BNG) (DEFRA, 2024). The England BNG metric calculates biodiversity ‘units’ before and after development by multiplying site size by a biodiversity score that is itself derived by multiplying a score for habitat distinctiveness by a score for habitat condition and a score for strategic significance. Habitat distinctiveness is determined by the rarity and conservation significance of the habitat type, which is subsequently multiplied by a value for habitat condition, representing the ecological quality of the habitat in question(DEFRA, 2024). This is then multiplied by a value for strategic significance which assigns a higher score to sites which are fulfilling the objectives of the relevant Local Nature Recovery Strategy (LNRS) (DEFRA, 2024).

Here, we test whether scores for grassland and arable sites generated from England’s BNG metric correlate with measures of ground invertebrate biodiversity based on samples collected using pitfall traps. Ground invertebrates are taxonomically and functionally diverse, with short generation times (Gerlach et al., 2013; Perner & Malt, 2003) and limited mobility compared to more frequently-studied groups such as birds, mammals or butterflies. These factors make them an especially useful focal group, as their measured richness, abundance and species composition will respond at the fine spatial scales relevant to the application of area-condition metrics.

Not only does the BNG Metric in England underpin a major national policy goal, but it is highly influential worldwide, with derivatives emerging globally including in Singapore, Sweden, the USA, Saudi Arabia, the Netherlands, and India (Supplementary Material 1) (Ecogain, 2023; Miller, 2024; Natural England, 2024; Ramboll, 2024). Therefore, its performance has direct relevance to the broader question of how effective metrics based upon habitat distinctiveness and condition are at representing biodiversity.

## Methods

To test the correlation between BNG metric score and measures of invertebrate biodiversity we sampled a range of habitat types representing combinations of habitat distinctiveness and condition scores. We focused on grassland habitat parcels in agricultural landholdings, as well as areas managed for wildlife conservation. This focus was motivated by a study of early BNG-adopting councils, which found that grasslands, in particular ‘modified grasslands’ and ‘other neutral grasslands’, are the most common habitats in BNG metric calculations (zu Ermgassen et al., 2021).

### Site selection and survey

In 2022, 14 habitat parcels (with habitat parcel defined as an area of the same habitat type and condition) were sampled from five landholdings in Oxfordshire and Buckinghamshire. Under the UK Habitat Classification system (UKHab) (UKHab, 2023), these were assessed to be five ‘modified grassland’ parcels, seven ‘other neutral grassland’ parcels, and two ‘arable field margin’ parcels. The 2022 parcels ranged from 0.6 ha to 12.3ha, with a mean parcel size of 4.0ha. In 2023, an additional set of 22 habitat parcels were investigated from five landholdings in Oxfordshire, Gloucestershire, and Buckinghamshire. These were assessed under UKHab to be two ‘temporary grass and clover ley’, seven ‘modified grassland’, seven ‘other neutral grassland’, four ‘lowland meadow’, and three ‘wood pasture and parkland’ habitat parcels. In 2023, habitat parcels ranged from 0.2ha to 8.7ha with a mean area of 2.4ha.

In both years, statutory biodiversity metric scores were generated for each habitat parcel using standard protocols for calculating BNG metric scores: (1) desk study to identify any statutory or non-statutory site designations using the DEFRA Magic Map (Natural England, 2023); (2) area delineation in the field; (3) floral survey; (4) habitat classification using UKHab 2.0 (UKHab, 2023); (5) metric condition assessment appropriate for habitat type (DEFRA, 2024); (6) calculation of biodiversity units using the statutory biodiversity metric (DEFRA, 2024). For the floral survey, between 5-10 1 m^2^ quadrats were used, increasing with the size and heterogeneity of the habitat parcel. These were placed to ensure that the quadrat configuration captured any site heterogeneity (e.g., soil moisture and slope). The floral survey was used to derive the UKHab type and subsequently the distinctiveness category of each habitat parcel. Habitat condition was assessed using the quadrat data and by extensive site walkovers to identify the extent of features that affect habitat condition (e.g., machinery damage or scrub encroachment) as laid out in the statutory biodiversity metric condition assessment methodology (DEFRA, 2024)

### Invertebrate surveys

Pitfall traps were used to sample ground invertebrate species richness and abundance, with the design intended to be easily standardised, minimise toxicity to non-target organisms, and to avoid creating an attractant to particular invertebrate groups (Woodcock, 2008). In the results we refer to invertebrate abundance, but it is important to note that pitfall trapping reflects invertebrate activity-abundance rather than abundance *per se*. In 2022, the pitfall traps used a twin plastic cup (width: 94mm, height: 135mm) containing 90ml of water, 10ml of propylene glycol, and <1ml of detergent. Galvanised chicken wire (mesh size: 35mm) was placed above the trap and flush with the ground surface to prevent vertebrate bycatch, and a black corrugated plastic rain cover (diameter: 150mm) was positioned 30mm above each trap, secured with tent pegs. Collected specimens were transferred to ethanol. In 2023, the trap design was modified by installing X-shaped rigid foam guidance barriers 650mm in length and 70mm in height, with the modified trap design increasing the trap radius and volume of catches (Boetzl et al., 2018). In each habitat parcel, the number of traps per parcel varied between three and nine, increasing with size and heterogeneity of the habitat parcel. As with the quadrats, the traps were placed to ensure that the configuration captured site heterogeneity.

In 2022, the traps ran for 4 weeks during August and September. Pitfall traps were active for approximately two weeks and then emptied and reset for another two weeks. In 2023, the traps were set for 1 week of each month in May, June, and August. The shorter timescale was appropriate because of higher catches arising from using guidance barriers and to prevent DNA degradation. Samples that were interfered with by vertebrates or otherwise damaged were removed from the dataset. In 2022, a total of 84 pitfall catches were collected across all sites, with each catch representing 28 trap-days, totalling 2352 trapping days. In 2023, 252 pitfall catches were collected across all sites, with each catch representing 7 trap-days, totalling 1764 trapping days.

### Invertebrate sorting and identification

In both years, the mean total invertebrate abundance of each parcel was taken by counting the number of individuals across major taxonomic groups: beetles (Coleoptera), spiders (Araneae), woodlice (Oniscidea), centipedes (Chilopoda), millipedes (Diplopoda), worms (Annelida), slugs and snails (Stylommatophora), and harvestmen (Opiliones). In 2022, ground beetles (Carabidae) were identified to species using Luff (2007). In 2023, to examine invertebrate species richness trends in more depth, samples were subjected to DNA metabarcoding by a commercial facility (NatureMetrics Ltd., Guildford, UK) using standard protocols (Supplementary Material 2). Due to resource constraints, the pitfall traps from each habitat parcel for each individual replicate were pooled. This decision was taken rather than pooling the temporal replicates for each individual pitfall trap so that seasonal variation could be examined. This meant that variation in DNA metabarcoding outcome within each habitat parcel between pitfall traps could not be examined, but we were able to retain this resolution for the abundance data.

### Data analysis

Data analysis and plotting was conducted in R using the packages ggplot2, vegan, dplyr, tidyverse, MASS, lme4, lmerTest, DHARMa, scatterplot3d, and iNEXT (Bates et al., 2015; Dixon, 2003; Hartig, 2022; Hsieh et al., 2016; Kuznetsova et al., 2017; Ligges & Maechler, 2003; R Core Team, 2024; Venables & Ripley, 2002; Wickham, 2016; Wickham et al., 2019, 2023). Generalised Linear Mixed Models (GLMMs) were fitted to test the relationship between distinctiveness*condition score and invertebrate abundance and species richness. Landholding was included as a random effect in all models to account for the spatial structure of the habitat parcels in the dataset. A negative binomial error distribution was used as it greatly improved model diagnostics relative to Poisson errors, with the ratio of residual deviance to degrees of freedom close to 1 in all cases. Degrees of freedom were extracted using the Welch– Satterthwaite equation in the package lmerTest (Kuznetsova et al., 2017). We conducted post-hoc power analysis using the package simr (Green & MacLeod, 2016). We also repeated the analyses in a Bayesian framework, using the same negative binomial mixed-effects model structure, and estimated model slopes and credibility intervals using the brms package (Bürkner, 2017). Both analyses are presented in Supplementary Table 3.

### 2022 and 2023 – Invertebrate abundance data

GLMMs were fitted to test the relationship between distinctiveness*condition score of habitat parcels and mean total invertebrate abundance, as well as carabid beetle and spider abundance.

### 2022 – Carabid species richness data

Carabid species richness was rarefied using coverage-based rarefaction to account for variation in sampling intensity and inventory completeness among parcels (Hsieh et al., 2016). Richness was rarefied to the minimum sample coverage of all area habitats (0.8575). The mean and confidence intervals were generated using 50,000 bootstraps. As above, GLMMs were fitted to test the relationship between rarefied carabid species richness and distinctiveness*condition scoring of habitat parcels.

### 2023 – Species richness data

DNA metabarcoding generated 689 Operational Taxonomic Units (OTUs) across all samples. OTUs from taxonomic groups not explicitly targeted by pitfall traps (non-ant Hymenoptera, Orthoptera, Lepidoptera, Mecoptera, Diptera, Nematoda, Plecoptera, Siphonaptera and Thysanoptera) were removed from the dataset, as these groups are considered to be by-catch, resulting in 417 OTUs from the target ground-dwelling invertebrate taxa. For analyses of OTU richness, OTUs which occurred only once in the dataset were removed, leaving 263 OTUs. However, only 50.5% of OTUs were resolved to a species-level identification. Given that multiple OTUs can have the same species-level identification, we also analysed species richness alongside OTU richness. To do this, we removed OTUs that did not have a species-level identification and amalgamated multiple OTUs that had the same species-level identification. This left 145 OTUs which were used to analyse total ground invertebrate species richness and the species richness of the main taxonomic groups.

Both OTU and species richness were rarefied to account for differences in sampling intensity using iNEXT (Hsieh et al., 2016). Richness was rarefied to a minimum sampling coverage of 0.55 to account for variation in inventory completeness and samples which fell below this threshold were excluded from the analysis. As above, GLMMs were used to test the relationship between total rarefied invertebrate OTU and species richness and distinctiveness*condition scores.

Due to zero inflation in the data and small datasets for individual taxonomic groups, sample coverage rarefaction was not possible for the species richness of individual taxonomic groups. Instead, the raw species richness value was used, excluding samples with <8 pitfall traps in total, and the number of pitfalls per parcel per season (between 8 and 15) was used as a covariate in the models to account for differences in sampling intensity. This was done for carabid beetle (Carabidae), rove beetle (Staphylinidae), spider (Araneae), slug and snail (Stylommatophora), true bug (Hemiptera), woodlouse (Oniscidea), and springtail (Collembola) species richness. Sampling intensity was not a significant predictor of species richness for any group. GLMMs were fitted to test the relationship between raw species richness and distinctiveness*condition scoring of habitat parcels, with landholding and number of pitfall traps per parcel as random effects.

### 2023 – Community Composition

To compare community composition across sites we used the OTUs from the DNA metabarcoding dataset which were identified to the family-level within our target pitfall taxa for May, June, and August (Hemprich-Bennett et al., 2021). This included 381 OTUs from 78 families. The species-level dataset was too sparse to produce a meaningful output from this analysis. Using numbers of family-level OTUs reduced the zero-inflation of the data and allowed Nonmetric Multi-dimensional Scaling (NMDS) clustering analysis. We used the number of family-level OTUs per sample and calculated a distance matrix using Bray-Curtis distance in the R package ‘vegan’ with 1000 permutations on 3 dimensions until the least-stress solution had been calculated (Dixon, 2003).

### Species of conservation concern

We deemed species to be of conservation concern if they were included as species of principal importance in England (England biodiversity list) or as species assessed as nationally scarce from relevant taxonomic assessments (DEFRA, 2022; Natural England, 2014, 2016).

## Results

### Invertebrate abundance and distinctiveness*condition score

We found no evidence for a relationship between a habitat parcel’s distinctiveness*condition score calculated using the BNG metric and the abundance of ground invertebrates, for grassland and arable habitat parcels surveyed in August and September 2022 (Fig 1a. z_10_= −0.86, p=0.39). In 2023, using a more efficient pitfall trap design and surveying a different set of habitat parcels encompassing a wider range of grassland and arable habitat types, there was similarly no significant relationship between distinctiveness*condition score and mean total ground invertebrate abundance (May: Fig 1b. z_14_= 1.86, p=0.06, June: Fig 1c. z_17_=0.99, p=0.32, August: Fig 1d. z_15_=0.16, p=0.88). There was also no significant relationship in either year between the abundance of the dominant groups in the samples (carabids and spiders) and distinctiveness*condition scores (p>0.1).

**Figure 1.**
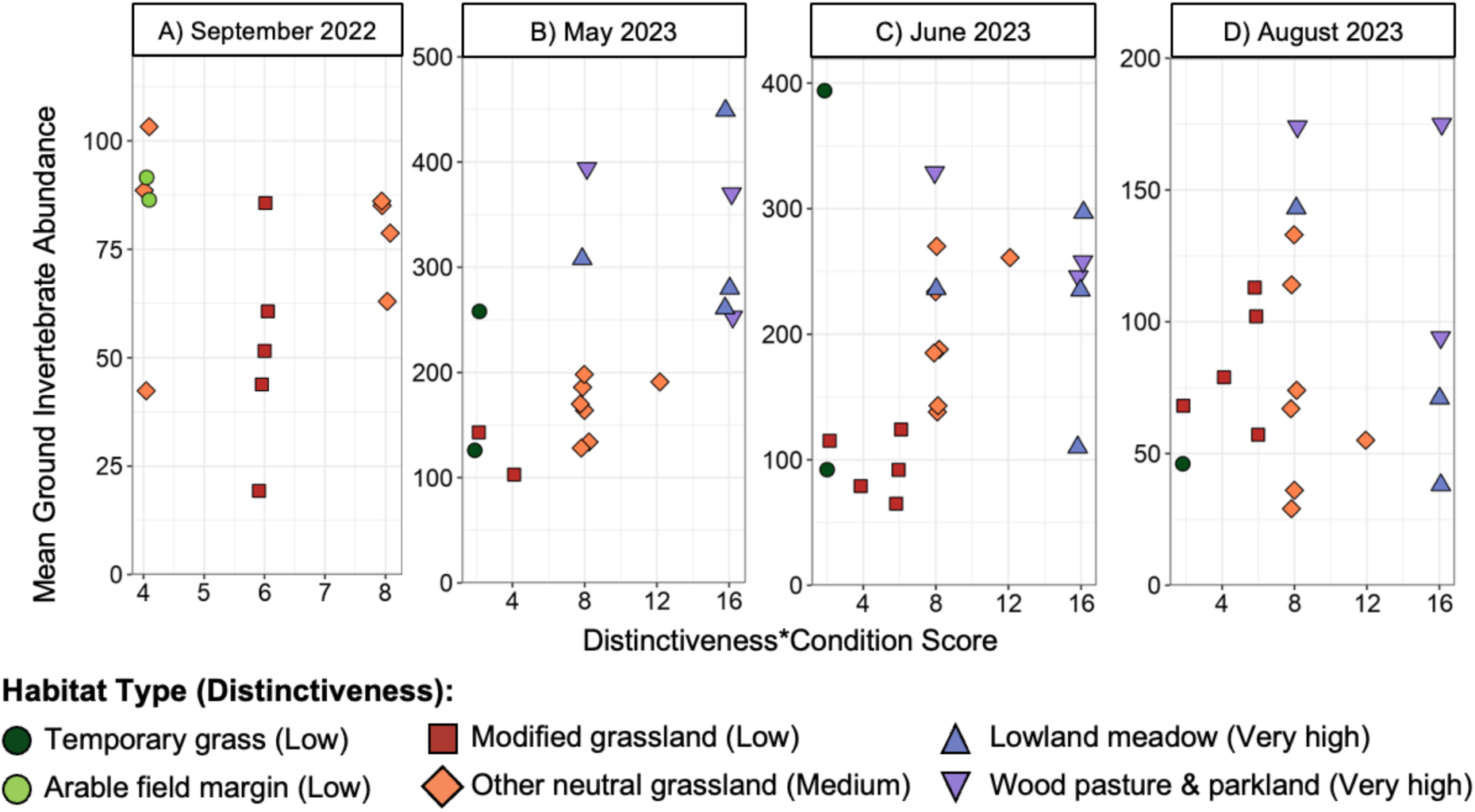
Mean ground invertebrate abundance and distinctiveness*condition scores over four weeks in 2022 (A), and three one-week periods across three months in 2023 (B,C,D).

### Species richness and distinctiveness*condition score

In 2022 there was no significant relationship between rarefied carabid beetle richness and distinctiveness*condition score (Fig 2a. z_10_= −1.28, p=0.20). In 2023, there was no significant relationship between total rarefied ground invertebrate species richness quantified by DNA metabarcoding and distinctiveness*condition score (Fig 2b. z_15_= 0.53, p=0.59). There was also no significant relationship between total rarefied Operational Taxonomic Unit (OTU) richness and distinctiveness*condition score (z_15_=0.54, p=0.60).

**Figure 2.**
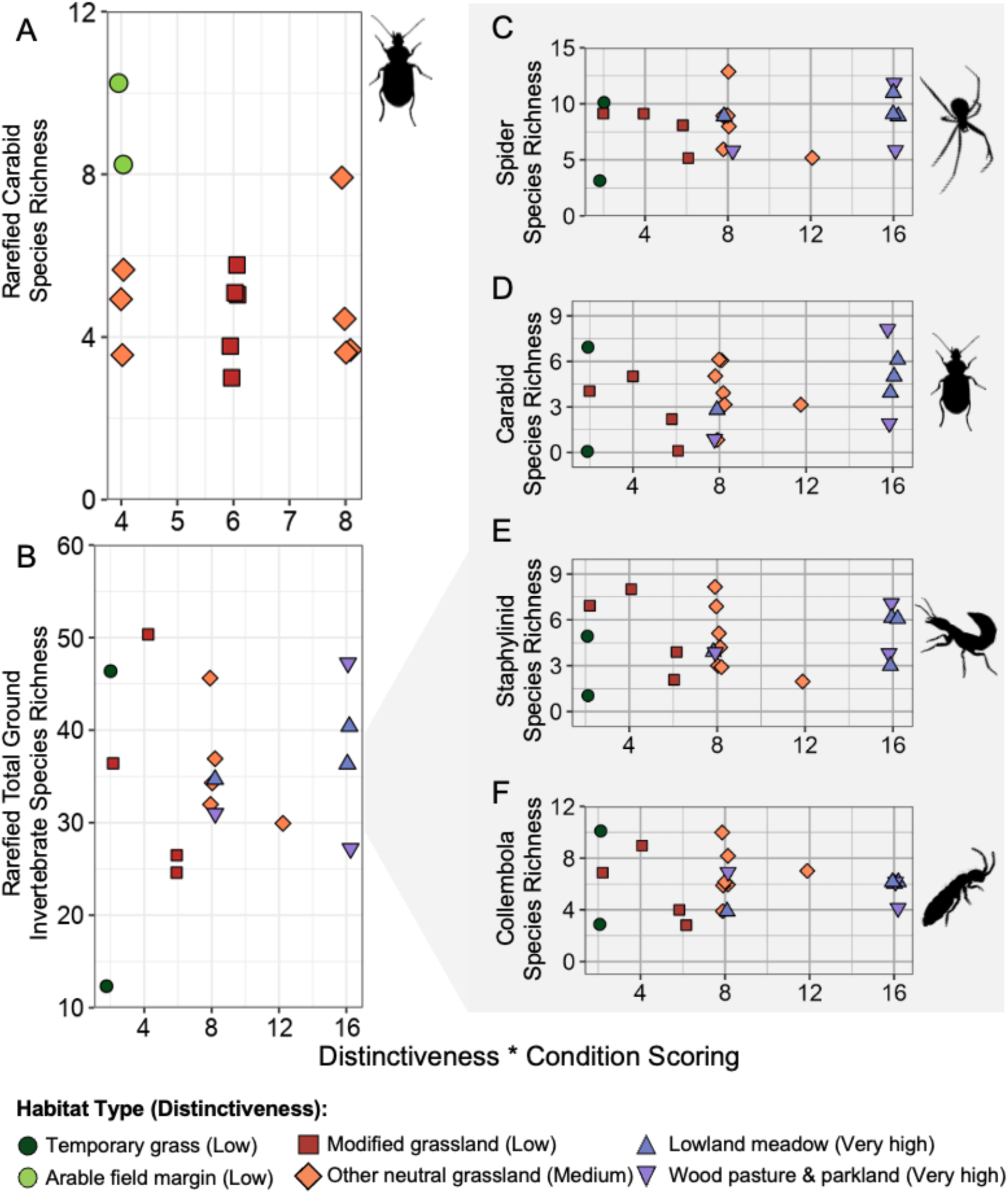
Ground invertebrate species richness and distinctiveness*condition scores for carabid beetles (A), total ground invertebrate species richness (B), spiders (C), carabids (D), staphylinids (E) and Collembola (F). Data for (A) are from 2022 sampling, and (B to D) from 2023.

Focusing on individual taxonomic groups, there were no evidence of relationships between distinctiveness*condition scores and species richness quantified using DNA metabarcoding for spiders (Fig 2c. z_15_=0.82, p=0.41), carabid beetles (Fig 2d. z_15_=1.19, p=0.23), rove beetles (Fig 2e. z_15_=0.15, p=0.88), Collembola (Fig 2f. z_15_ =-0.57, p=0.56) or Stylommatophora (z_15_= 0.39, p=0.69).

### Community composition

A 3D Nonmetric multidimensional scaling (NMDS) analysis of the number of family-level OTUs derived from DNA metabarcoding in each habitat parcel shows the clustering of invertebrate communities by habitat type in May, June, and August 2023 (Fig 3a-c). The ordination revealed almost complete overlap of communities drawn from habitats with differing distinctiveness scores under the BNG metric. In August 2023, there is some evidence of separation between lower and higher distinctiveness habitats, although not between habitat types within those distinctiveness groupings.

**Figure 3.**
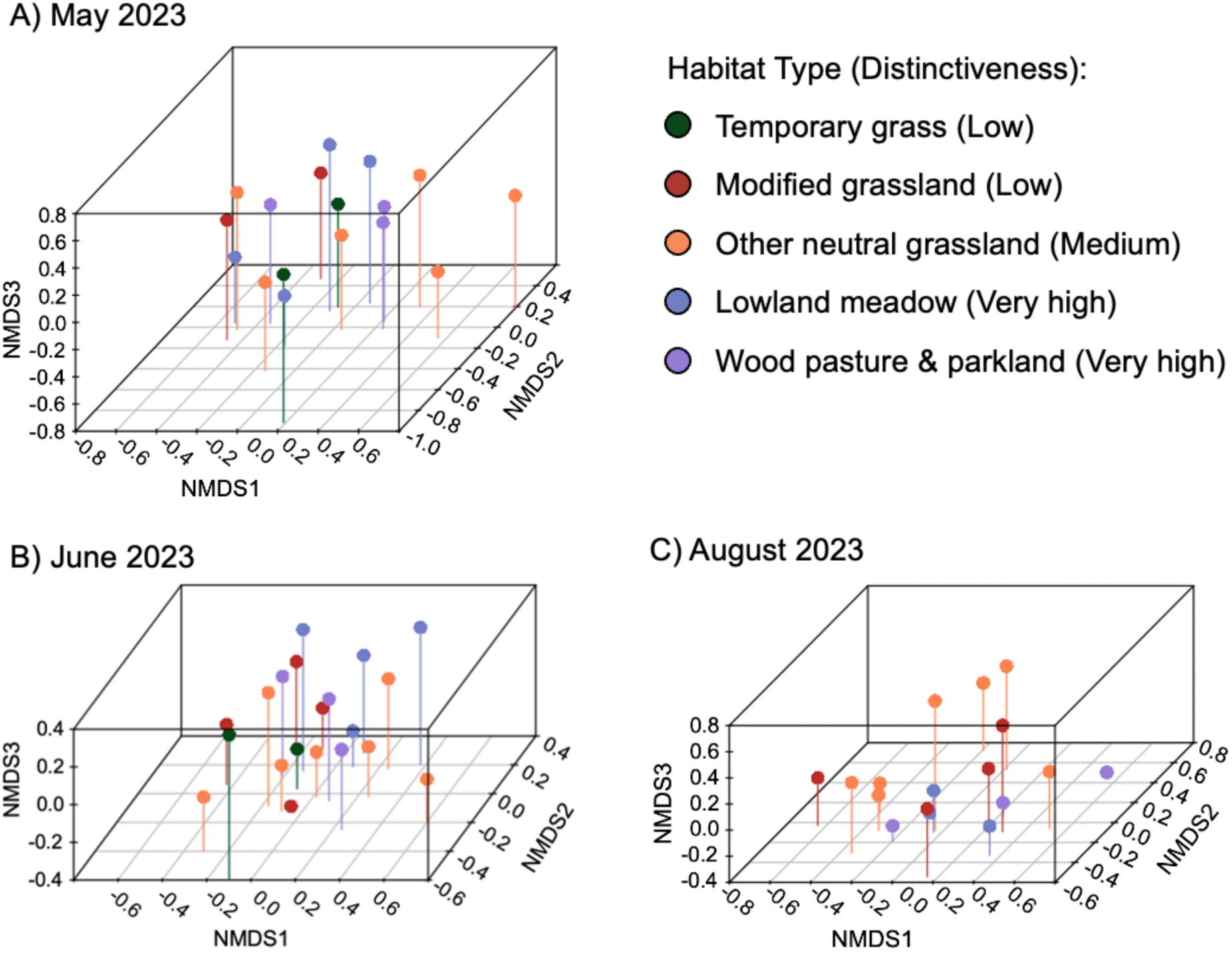
Nonmetric multidimensional scaling of family-level Operational Taxonomic Units (OTUs) from 2023. The least stress solution was 0.09 for May (A), 0.12 for June (B), 0.13 for August (C).

### Species of conservation concern

We recorded 13 occurrences of 7 species which are nationally scarce in the United Kingdom, including Biodiversity Action Plan (UKBAP) species. These were found with indistinguishable frequency across a range of habitat parcels, from ‘low’ to ‘very high distinctiveness’ and ‘poor’ to ‘good condition’ (Supplementary material 4).

## Discussion

Combined area-condition biodiversity metrics are widely applied proxies for evidencing improvements to biodiversity, and their use is proliferating rapidly as demand for easy-to-use biodiversity quantification methods rises to satisfy many policy and business objectives (Table 1). Here we fail to find evidence that a prominent combined area-condition metric (the BNG metric) is related to the abundance and species richness of ground invertebrates. If use of these metrics result in trades of large, low-scoring sites for smaller, high-scoring sites (as observed in practice: Rampling et al. 2023) then wider biodiversity benefits may not be achieved, or even counter-incentivised, resulting in accelerated biodiversity loss. In England, harm to high-scoring sites such as lowland meadows is strongly disincentivised, and where it does occur it requires bespoke compensation to be agreed with local authorities. This is not the case for lower-scoring sites, and if the metric is the only measure of biodiversity value considered, our results therefore imply regular undervaluing of sites of high value for invertebrates, which is likely also to be an issue for other components of biodiversity.

**Table 1.** Components of the Statutory Biodiversity Metric used to calculate baseline biodiversity value.

| Term | Definition | Application |
| --- | --- | --- |
| **Baseline metric components** (*size \* distinctiveness \* condition \* strategic significance*) |  |  |
| *Size* | Area or length of a given habitat feature | Multiplies metric score by area (hectares) or length (kilometres) |
| *Habitat distinctiveness* | A measure of the conservation value of a given habitat type, based on its rarity, existing protections, and its importance for species richness and specialist species. | Multiplies metric score by 2 (low), 4 (medium), 6 (high), or 8 (very high) |
| *Habitat condition* | A measure of the ecological condition of a habitat relative to its optimum state | Multiplies metric score by 1 (poor), 2 (moderate), 3 (good) |
| *Strategic significance* | A measure of whether a habitat delivers on local nature recovery priorities | Multiplies metric score by 1 (low), 1.10 (medium), 1.15 (high) |

If, as we found here, combined area-condition metrics are not reliable proxies for wider biodiversity value, sites with high faunal biodiversity value but low metric scores will be undervalued in offsetting calculations. In this scenario, a low baseline metric score would mean fewer offsets would be required than would be necessary to balance the environmental damage caused by a development requiring ecological compensation. For voluntary biodiversity credit initiatives, an inaccurate proxy metric would lead to investment in credits with uncertain or even overinflated biodiversity benefits. If such biodiversity credits become central to how businesses claim to achieve ‘nature-positive’ objectives (i.e. claiming they deliver more improvements to biodiversity than damage) or address their nature-related risks under prevailing Corporate Sustainability Reporting frameworks and policies (e.g. the Taskforce for Nature-relate

Financial Disclosures or the EU Corporate Sustainability Reporting Directive), using a metric that doesn’t reliably capture faunal diversity risks creating the illusion of biodiversity gains (Maron et al., 2024). Overestimating the biodiversity benefits delivered in these systems could have negative repercussions. In the case of the voluntary carbon market, evidence that carbon offset projects were not delivering as much gain as promised (West et al., 2023) led to a 56% reduction in the volume of the voluntary carbon market the following year (Forest Trends Ecosystem Marketplace 2024). Robust biodiversity metrics are essential to avoid overpromising and underdelivering, to prevent investors losing confidence in funding biodiversity conservation through biodiversity credit systems (Swinfield et al., 2024). Metrics such as the BNG metric seek to deal with this risk by incorporating negative multipliers, which require the loss of high value habitat to be replaced with a much larger area of the same habitat (Bull et al., 2013). However, these multipliers require careful modelling and design to ensure that they are set at the appropriate level to achieve that goal (Bull et al., 2017; Laitila et al., 2014; Moilanen et al., 2009).

Our results are also concerning given evidence of declining populations of ground invertebrate species (Brooks et al., 2012; Ewald et al., 2024; Mancini et al., 2023), their contribution to important ecosystem functions such as nutrient cycling, pest control, and decomposition (Dainese et al., 2019; Nichols et al., 2008; Seibold et al., 2021), and their role as food resources for taxa such as farmland birds (Holland et al., 2006). Recent analyses suggest that the BNG metric is also a poor proxy for birds and other vertebrates of conservation concern (Hawkins et al., 2022; Marshall et al., 2024), although the higher mobility and larger territories and home ranges of most vertebrates make it less likely that patterns in their abundance and diversity align with BNG metric scores in fine-grained, heterogeneous landscapes such as much of lowland England, where parcel sizes or habitat units for which BNG scores are calculated are small. Given the importance of the ecosystem functions fulfilled by ground invertebrates and their significance in food webs, optimising combined area-condition metrics to better reflect their habitat requirements could trigger better habitat protection and compensation for other taxa. Indeed, certain groups of ground invertebrates, such as ground beetles, are good proxies for the diversity of other invertebrates, vertebrates, and plants (Schuldt & Assmann, 2010). Conversely, optimisation of metrics for vertebrate requirements may possibly result in outcomes antithetical to invertebrate biodiversity, where conservation planning assumes vertebrates will have an ‘umbrella effect’, without accounting for patterns of invertebrate richness and distribution (Oberprieler et al., 2019).

Like many other combined area-condition metrics, the BNG metric takes a habitat-based approach (Treweek et al., 2010). However, our results indicate that the habitat type classification used in the metric does not predict invertebrate biodiversity, with high variability between sites of the same habitat type. The BNG metric is underpinned by the UK Habitat Classification (UKHab) system, and each habitat type within the metric constitutes an umbrella for a range of different sub-communities (UKHab, 2023). This could explain some of the variation we observed between sites classified as the same habitat type but does not account for the higher invertebrate species richness and abundance observed in some ‘low’ and ‘medium distinctiveness’ grassland sites compared to ‘very high distinctiveness’ sites. Our study focused on a subset of all the habitats in BNG (grasslands), because alongside cropland these habitats are by far most consequential and frequently occurring in BNG assessments across the country. In focusing on a narrow suite of habitats, we were able to use our intense sampling effort to explore variation within habitat types as well as between habitat types. We found no consistent patterns in the relationship between invertebrate communities and habitat type or condition, with our models finding very small effect sizes. Our small sample size does not lend enough statistical power to give us confidence in accurately detecting such a small effect size. However, given that the BNG metric operates on a site-by-site level, the magnitude of difference between levels in the metric should be sufficiently great to differentiate the biodiversity value of even a small selection of sites. Here, we see that this is not the case for the relationship between the BNG metric and invertebrate biodiversity. This highlights the significance of ground truthing and piloting in metric design to ensure that the differentiation in the scoring system is ecologically meaningful.

The condition assessment applied to grasslands also risks exacerbating the mismatch between metric scores and invertebrate biodiversity by devaluing sites with habitat features of value to invertebrates (Duffus et al., 2024). Sites with more than 5% cover of scattered scrub, bare ground, and plant species ‘indicative of sub-optimal condition’ are assigned a lower score, thereby providing an incentive for the removal of these features when they arise (DEFRA, 2024). This simplified scoring does not recognise the value of bare ground and scrub for creating heterogenous sites which support unique ground invertebrate assemblages (Key, 2000; Lyons et al., 2018). While vegetation-based features simplify monitoring and capture some useful elements of biodiversity, it is important that the features chosen are relevant to a wider range of flora and fauna.

Refinement of the BNG metric habitat classification and condition assessment may still not capture important dimensions explaining variations in biodiversity among sites. Landscape-level factors are important determinants of a site’s biodiversity, not only for invertebrates, but also for groups such as insect pollinators, bats, and birds (Le Provost et al., 2021). Landscape-level factors such as connectivity, which is particularly important for population persistence of less mobile, more specialised taxa (Wamser et al., 2012), are not currently captured by the metric. Site age is also a consideration, given the long time-scales over which some invertebrates take to colonise new sites (Woodcock & McDonald, 2010). Developing a way to incorporate site age and connectivity could ensure that older, well-connected sites achieve higher scores, reflecting a higher likelihood of well-established invertebrate communities. For example, the BNG metric uses a measure of ‘strategic significance’ which assigns a higher score to post-development habitats which contribute to the delivery of local priorities for nature recovery (Department for Environment, Food & Rural Affairs, 2023). However, this does not recognise the connectivity value of sites pre-development, such as a low scoring grassland that connects high scoring grasslands. Previously, metric 2.0 incorporated a connectivity multiplier, but it was removed because users found it challenging to implement, highlighting the importance of retaining practicality (Natural England, 2020).

Metrics based upon habitat area, type, and condition could be refined to better represent faunal diversity by: using a robust appropriately fine-grained habitat classification system; selecting indicators of site quality that reflect the habitat requirements of a range of faunal species; and incorporating landscape-scale metrics such as connectivity. However, in the absence of evidence that combined area-condition metrics are robust proxies for faunal diversity, metrics should be supplemented with on-the-ground species data collection to understand the biodiversity present on sites under consideration for either development or conservation action. This is especially pertinent where conservation evaluations or prioritisation are occurring. We also found large differences in community composition in different months within a single season (Fig 3a-c), demonstrating the need for repeated data collection to account for seasonal variation, with interannual variation (not quantified here) also likely. Using species data alongside combined area-condition metrics would help identify sites with a low metric score which nonetheless support populations of species of conservation concern, or highly diverse or unique communities. However, in England a shift away from collecting species data is currently proposed in an effort to streamline the planning process (Planning and Infrastructure Bill, 2025). Our results demonstrate the importance of species data in identifying site value and likely impacts on fauna, and enabling more nuanced decision-making which reflects the true biodiversity value of sites.

Amid high demand for easy-to-use biodiversity proxies, combined area-condition biodiversity metrics have become widely adopted tools for a range of biodiversity monitoring applications including offsetting schemes and biodiversity credit calculations. Once a particular metric becomes established, it is likely to be emulated by others (as is happening with the England BNG metric). This is good, if lessons from the implementation of a metric are used to improve it and to inform the development of other metrics. Our data demonstrate that area-condition metrics can be a poor proxy for invertebrate diversity sampled using pitfall traps, in terms of abundance, richness, community composition, and the occurrence of species of conservation concern. Combined area-condition metrics could be improved by refining indicators of ecological quality to better reflect habitat features required by a range of species, including invertebrates. However, without accompanying data on the occurrence of these species, uncertainty will remain about the full biodiversity benefits of interventions designed using combined area-condition metrics. Ultimately, this leads to the risk that development will occur on sites of high biodiversity value, that losses to biodiversity will not be fully offset, or that the biodiversity benefits of biodiversity credits will be overestimated.

## Author Contribution Statement

O.T.L., E.J.M.G., R.G., S.O.S.E.z.E., N.E.D., and T.B.A. conceptualised the study. N.E.D., T.B.A., D.C., B.S., and N.T., collected the data. N.E.D. and T.B.A. analysed the data and produced the figures. N.E.D. and T.B.A. wrote the initial manuscript. O.T.L., R.G., S.O.S.E.z.E., J.W.B., E.J.M.G., N.E.D., and T.B.A., contributed to subsequent iterations of the manuscript and all authors approved the final version for submission.

## Data and Code Availability

Data and code from this paper can be shared by lead author on request and will be deposited and accessible on Dryad prior to the paper’s publication date.

## Acknowledgements

The authors would like to thank the Agile sprint team for their support on this project, including Prue Addison and Julia Baker for their advice on fieldwork and the use of the metric, and Isobel Hawkins, Dom Meeks, and Lucy Jones for their assistance with data collection. We also thank Community Ecology Research Oxford (CERO) for their support, including Dave Hemprich-Bennett and Chris Terry for their valuable advice on the data analysis. Finally, we would like to thank all the landowners for allowing us the use of their sites for this study.

