## Supplementary Materials for "A globally influential area-condition metric is a poor proxy for invertebrate biodiversity"

**Supplementary Material**

**Supplementary Material 1: Global derivatives of England’s statutory biodiversity metric**

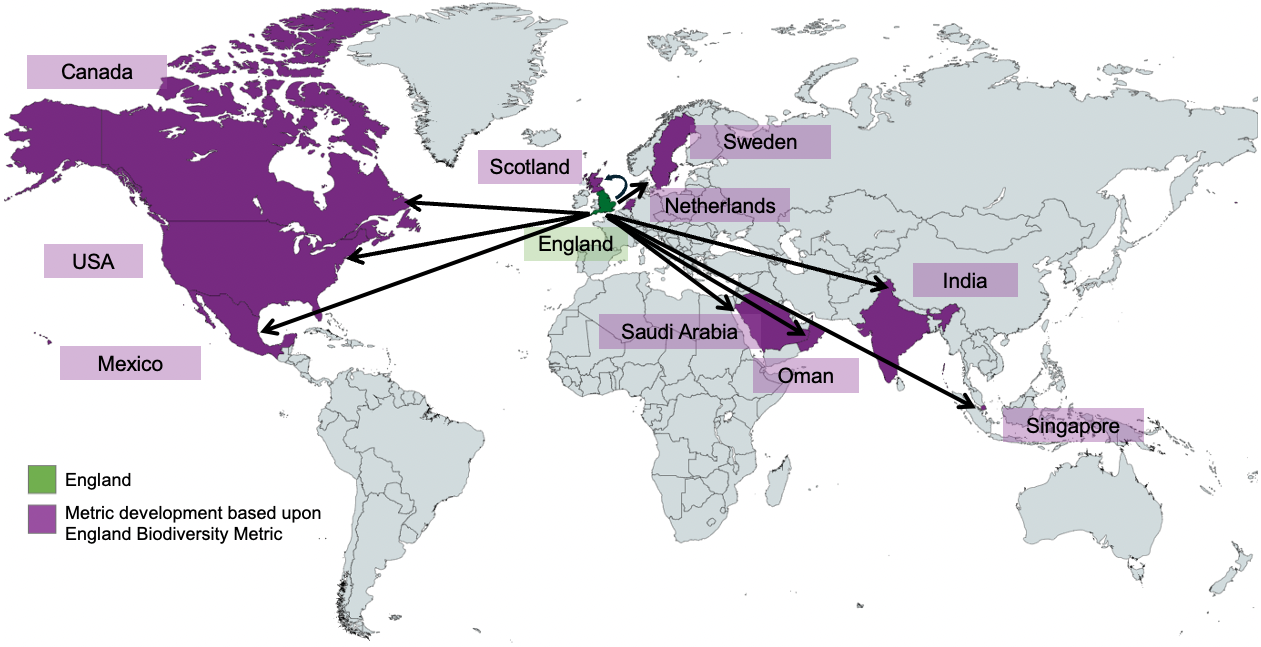

SI Fig 1: Countries for which an adaptation of the England statutory biodiversity metric is developed/in development.

**Supplementary Material 2: DNA metabarcoding process**

NatureMetrics used the following standard protocol for the DNA metabarcoding of samples in this study.

***Bulk Invertebrate Extraction (non-destructive)***

A dedicated clean room was used for sample processing, using class II biosafety cabinets and workstation decontamination before and after use with chemical disinfectant and UV irradiation. Samples were rinsed with ethanol and dried at 30°C overnight. A lysis buffer (modified from Ivanova et al., 2006) was then added to the samples which were incubated at 56°C for up to 4 hours, depending on the invertebrate volume. A DNeasy Blood and Tissue Kite (Qiagen) was used to process an aliquot of the lysate. A negative control of lysis buffer to monitor exogenous DNA contamination was also processed with each batch of samples. DNA concentration was measured to determine extraction yields using a Qubit fluorometer with Qubit dsDNA broad range assay kit (Thermo Fisher Scientific).

***DNA Amplification***

Each sample and extraction blank was amplified using a two-step PCR process. Tails were added to the 5’ end of taxon-specific primers to complement adapter and index primer sequences downstream. Hot Start DNA polymerase was used for amplification using forward primer mlCOIintF-XT 5 and jgHCO2198, targeting the mt-COI gene for insects. Proprietary synthetic sequences that do not match known biological records and PCR-grade water were used as positive and negative controls, respectively. These controls were included with every PCR plate to verify the amplification performance. PCR amplification success was confirmed visually by gel electrophoresis.

***Library Preparation and Sequencing***

First round PCR replicates that were amplified successfully were pooled per sample and purified using MagBind TotalPure NGS magnetic beads (Omega Biotek). Then, from the purified amplicons, a sequencing library was prepared, using unique dual indexes, following Illumina’s 16S Metagenomic Sequencing Library Preparation protocol (Illumina, 2024). Indexed PCR products were purified, quantified, normalised, and pooled in equal volumes. An Illumina MiSeq system using a V3 600 cycle reagent kit was used to sequence the final library.

***Bioinformatics***

Sequences were demultiplexed with bcl2fastq and processed using a custom NatureMetrics eDNA analysis pipeline. USEARCH was used to merge the paired-end FASTQ reads generated for each sample (Edgar, 2010). Cutadapt was used to trim the forward and reverse primers from the merged sequences, with a length filter of 300-330bp (Martin, 2011). USEARCH was used for filtering the quality of sequences. Only sequences with an expected error rate of 0.01 per base and below and dereplicated by the sample were retained. This retained singletons to obtain zero-radius Operational Taxonomic Units (zOTUs). The unique sequences from all samples were denoised using UNOISE (Edgar, 2016).

For each zOTU, a consensus taxonomic assignment was made using sequence similarity searches against the NCBI nucleotide reference database SILVA (Quast et al., 2013; Yilmaz et al., 2014), BOLD (Altschul et al., 1990; Ratnasingham and Hebert, 2007; Camacho et al., 2009) and required hits to have a minimum e-score of 1e-20 and cover at least 90% of the query sentence. For all hits, the associated taxonomic identification was converted to match the GBIF taxonomic backbone.

Taxonomic assignments were made to the lowest possible level where matches were consistent, which the similarity thresholds set to a minimum of 98%, 95%, and 92% for species, genus, and higher-level assignments respectively. GBIF occurrence records were used to sense check the identifications made, checking for presence of species in the UK, and where necessary identifications were elevated to a higher taxonomic level (rgbif; Chamberlain et al., 2022).

zOTUs were clustered at 97% similarity with USEARCH to obtain OTUs. The dereplicated reads generated for each sample were mapped to the OTU representative sequences using USEARCH with an identity threshold of 97%. This mapping was used to create a table of OTU-by-sample. The OTU table was filtered to remove low abundance OTUs from each sample (<0.02% or <10 reads, whichever is greater threshold for the sample).

**Supplementary Material 3: Model power and coefficients from Bayesian framework**

Bayesian modelling was carried out in the brms package, using an identical model specification to the models presented in the manuscript: a generalised linear mixed-effect model with a binomial error structure and with landholding as a random effect. Uninformative prior distributions were set on all estimated coefficients, and posterior distributions were estimated via an adaptive NUTS sampler. In all cases, 6 chains were run with 1000 iterations of warmup and 50,000 sampling iterations. Following control optimisations (adapt_delta = 0.9999, step_size = 0.005, max_treedepth = 20), all models showed full convergence (Rhat = 1) and numerical stability except the model for 2022 total species richness, which is excluded here.

| **Model** | **Post-hoc power** | | | **Bayesian coefficient** | | | | |
| --- | --- | --- | --- | --- | --- | --- | --- | --- |
|  | **Power** | **Lower 95% CI** | **Upper 95% CI** | **Coefficient** | **Lower 95% CI** | **Upper 95% CI** | **Intercept** | **R^2^** |
| 2022 abundance | 21.5% | 14.33 | 31.39 | -0.06 | -0.21 | 0.1 | 4.65 | 0.53 |
| 2022 species richness | 33.% | 25.73 | 45.18 | N/A | N/A | N/A | N/A | N/A |
| 2023 May abundance | 52% | 43.74 | 64.02 | 0.02 | -0.01 | 0.06 | 5.19 | 0.67 |
| 2023 June abundance | 25% | 16.88 | 34.66 | 0.02 | -0.03 | 0.07 | 5.07 | 0.38 |
| 2023 August abundance | 9% | 4.20 | 16.40 | 0 | -0.06 | 0.07 | 4.47 | 0.34 |
| 2023 species richness | 6% | 1.10 | 9.93 | 0.01 | -0.03 | 0.04 | 3.46 | 0.17 |

**Supplementary Material 4: Table of species of conservation concern**

**Table 3.** Occurrence of species of conservation concern in the dataset

| **Species** | **Designation** | **Habitat Type** | **Distinctiveness** | **Condition** | **Distinctiveness * Condition score** |
| --- | --- | --- | --- | --- | --- |
| *Brachinus crepitans* (Linnaeus 1758) | Nationally scarce | Arable Field Margin | Medium | N/A | 4 |
| *Pterostichus anthracinus* (Illiger 1798) | Nationally scarce | Other Neutral Grassland | Medium | Moderate | 8 |
|  |  | Lowland Meadow | Very high | Moderate | 16 |
| *Meloe rugosus* Marsham 1802 | England biodiversity list | Other Neutral Grassland | Medium | Moderate | 8 |
| *Anaspis thoracica* (Linnaeus 1758) | Nationally scarce | Modified Grassland | Low | Good | 6 |
| *Liocranoeca striata* (Kulczynski 1882) | Nationally scarce | Other Neutral Grassland | Medium | Moderate | 8 |
| *Pardosa proxima* (Koch 1847) | Nationally scarce | Other Neutral Grassland | Medium | Moderate | 8 |
|  |  | Lowland Meadow | Very high | Moderate | 16 |
| *Carabus monilis* Fabricius 1792 | England biodiversity list | Lowland Meadow | Very high | Moderate | 16 |
|  |  | Lowland Meadow | Very high | Poor | 8 |
|  |  | Other Neutral Grassland | Medium | Moderate | 8 |
